# Altered A-type potassium channel function impairs dendritic spike initiation and temporammonic long-term potentiation in Fragile X syndrome

**DOI:** 10.1101/2021.01.06.425593

**Authors:** Gregory J. Ordemann, Christopher J. Apgar, Raymond A. Chitwood, Darrin H. Brager

## Abstract

Fragile X syndrome (FXS) is the leading monogenetic cause of cognitive impairment and autism spectrum disorder. Area CA1 of the hippocampus receives current information about the external world from the entorhinal cortex via the temporoammonic (TA) pathway. Given its role in learning and memory, it is surprising that little is known about TA long-term potentiation (TA-LTP) in FXS. We found that TA-LTP was impaired in *fmr1* KO mice. Furthermore, dendritic Ca^2+^ influx was smaller and dendritic spike threshold was depolarized in *fmr1* KO mice. Dendritic spike threshold and TA-LTP were restored by block of A-type K^+^ channels. The impairment of TA-LTP coupled with enhanced Schaffer collateral LTP may contribute to spatial memory alterations in FXS. Furthermore, as both of these LTP phenotypes are attributed to changes in A-type K^+^ channels in FXS, our findings provide a potential therapeutic target to treat cognitive impairments in FXS.

## Introduction

Fragile X syndrome (FXS), caused by transcriptional silencing of the *fmr1* gene and loss of Fragile X mental retardation protein (FMRP)^1^, is the leading monogenetic cause of autism and intellectual disability, affecting approximately 1 in 4000 males and 1 in 6000 females^2^. FMRP controls many neuronal proteins, including those involved in synaptic structure, function and plasticity, through translational regulation of target mRNAs^3^ and direct protein-protein interactions^4,5^. Given its high level of FMRP expression, well-defined circuitry, and relevance to learning and memory processes, the hippocampus has been critical to understanding synaptic changes in FXS^6-8^. The Schaffer collateral CA3 to CA1 synapse has been extensively studied in FXS^9^. By contrast, few studies have investigated the temporoammonic (TA) entorhinal cortex to CA1 synapses. Wahlstrom-Helgren and Kylachko found no difference in TA synaptic transmission using somatic whole cell recording^10^, while Booker et al. found that TA synaptic transmission was reduced in *fmr1* knock-out (KO) mice using extracellular field potential recording^11^. Neither of these studies, however, investigated TA long-term potentiation (LTP). Given the lack of LTP studies and the critical involvement of the TA pathway in the consolidation of long-term memory^12^ and its necessity for the generation of hippocampal CA1 place fields^13^, we asked whether TA-LTP is altered in FXS.

CA1 pyramidal neuron dendrites express Na^+^, K^+^, Ca^2+^, and h-channels which play crucial roles in dendritic integration and the induction of LTP. We previously showed that the functional expression of dendritic h-channels (I_h_) is higher in *fmr1* KO CA1 neurons^5,14^. We also demonstrated that the current carried by dendritic A-type K^+^ channels (I_KA_) is reduced in *fmr1* KO CA1 neurons^15^. These changes alter the local integrative properties and increase the backpropagation of action potentials respectively. While the effect of these changes in dendritic h-channels and I_KA_ on Schaffer collateral LTP were previously described^14,15^, the impact on TA-LTP remains unknown.

Using somatic and dendritic recording, we found that TA-LTP following theta-burst stimulation was impaired in *fmr1* KO mice. The lack of LTP was not due to the higher expression of h-channels as block of I_h_ with ZD7288 did not rescue LTP. Two photon imaging during bursts of TA stimulation revealed that dendritic Ca^2+^ signals were smaller in *fmr1* KO neurons. Although complex and pharmacologically isolated Ca^2+^ spikes recorded in the dendrites were not different, the threshold for fast dendritic spikes (dspikes) was more depolarized in *fmr1* KO CA1 pyramidal neurons. Application of extracellular Ba^2+^ or AmmTx3 to block I_KA_ rescued dendritic spike threshold and TA-LTP in *fmr1* KO CA1 pyramidal neurons, implicating A-type K^+^ channels.

## Results

### Long-term potentiation of TA synapses is impaired in fmr1 KO mice

We made somatic whole cell current clamp recordings from CA1 pyramidal neurons in wild type and *fmr1* KO male mice and recorded TA EPSPs before and after theta burst stimulation (TBS) to induce TA-LTP^16^ (Fig. 1a). In wild type neurons TA EPSP slope increased 30 minutes post TBS (Fig. 1b-c). By contrast, in *fmr1* KO CA1 neurons TA EPSP slope was not increased after TBS (Fig. 1b-c). The somatic depolarization during the TBS (area under the curve) was greater in wild type compared with *fmr1* KO CA1 pyramidal neurons (Fig. 1d). In agreement with previous work^10^, we found no difference in postsynaptic responsiveness to single TA stimuli Fig. 1e), baseline paired-pulse ratio (Fig. 1f) or temporal summation (Fig. 1g) between wild type and *fmr1* KO neurons. Paired pulse ratio was also not changed after TBS consistent with a postsynaptic locus of LTP (wild type: 2-way RM ANOVA: F(1,4)=3.53, p=0.13; *fmr1* KO: 2-way RM ANOVA: F(1,5)=3.98, p=0.1).

**Fig. 1:**
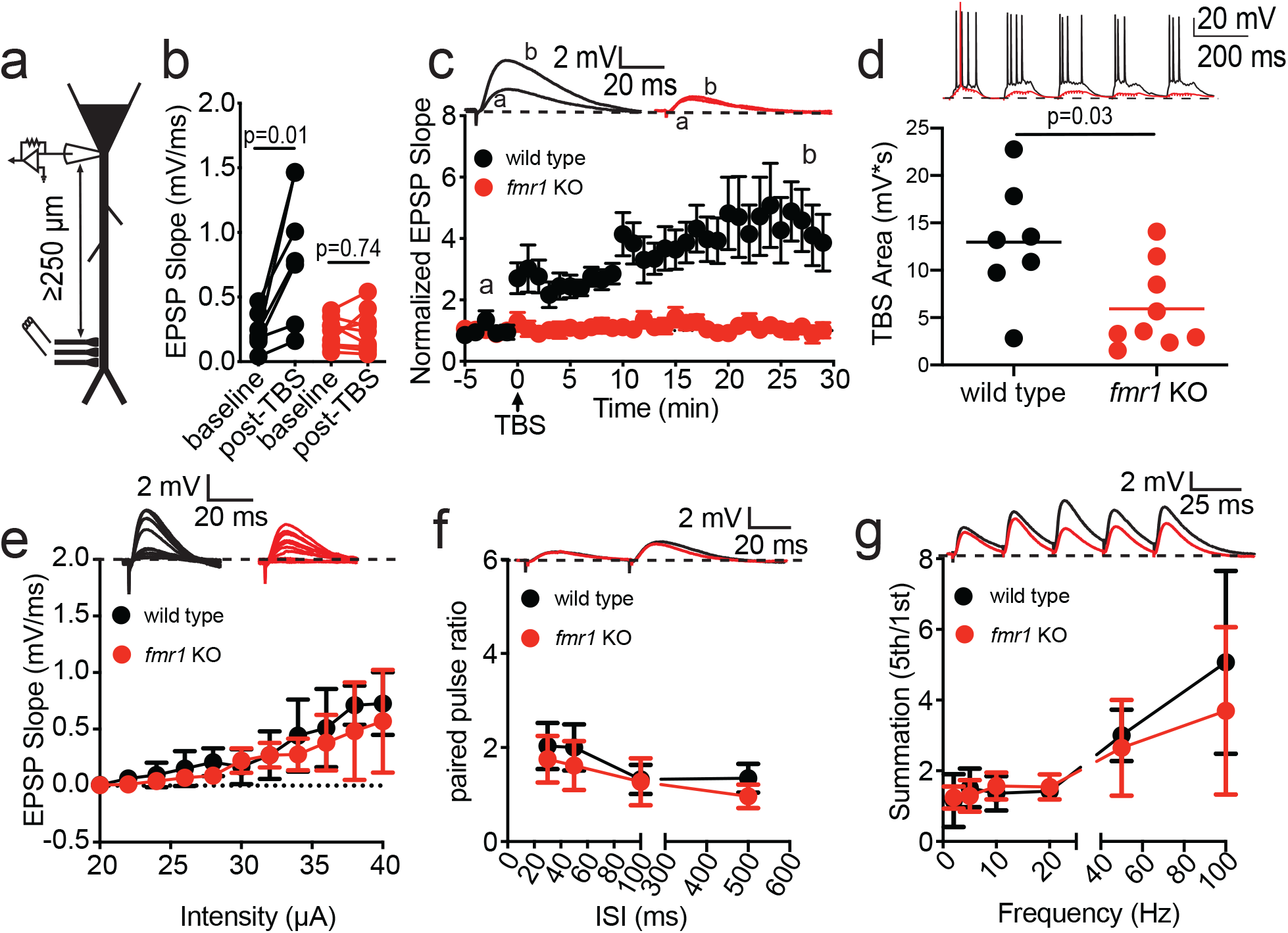
TA-LTP is impaired in fmr1 KO CA1 pyramidal neurons. **a,** Diagram of recording configuration for somatic EPSP measurements. **b,** EPSP slope is increased 30 minutes after TBS in wild type but not *fmr1* KO CA1 neurons; baseline: (mean of 5 minutes before TBS); post-TBS: (mean of 5 minutes at the end of recording); (wild type: baseline: 0.24 ± 0.054, post-TBS: 0.84 ± 0.19 mV/ms. paired t-test: t(6)=3.62, p=0.01, η^2^=0.39. *fmr1* KO: baseline: 0.23 ± 0.04. post-TBS: 0.24 ± 0.05 mV/ms. paired t-test: t(8)=0.35, p=0.74) **c,** Normalized change in EPSP slope for wild type and *fmr1* KO CA1 neurons. *Inset:* Representative EPSP traces from baseline (a) and post-TBS (b). **d,** Graph of area under the curve during TBS (unpaired t-test: t(10.37)=2.52, p=0.03, η^2^=0.31) **e,** Input output of TA inputs measuring EPSP slope (2-way RM ANOVA: F(1,7)=0.65, p=0.45). *Inset:* representative experiments from wild type and *fmr1* KO neurons. **f,** Paired pulse ratio as a function of interstimulus interval (ISI) (2-way RM ANOVA: F(1,9)=4.23, p=0.07). *Inset:* representative 50 ms ISI traces. **g,** Temporal summation of TA EPSPs as a function of frequency (2-way RM ANOVA: F(1,9)=0.55, p=0.48). *Inset*: representative 50 Hz traces.

We filled CA1 pyramidal neurons with neurobiotin during whole cell recording for post-hoc morphological reconstruction and analysis. We identified no differences in dendritic morphology between wild type and *fmr1* KO CA1 pyramidal neurons (Supp. Fig. 1a). Taken together, these data show that although basal TA synaptic transmission and CA1 neuron morphology are normal, TA-LTP is impaired in *fmr1* KO CA1 neurons.

### Block of I_h_ does not rescue TA-LTP in fmr1 KO neurons

The expression of h-channels in the distal dendrites of CA1 pyramidal neurons constrains TA inputs and LTP^16^. We previously showed that dendritic I_h_ is elevated in *fmr1* KO CA1 pyramidal neurons compared to wild type^5,14^. Higher I_h_ in *fmr1* KO CA1 neurons may reduce the effectiveness of TA synapses in the distal dendrites and impair LTP^17,18^. To test this hypothesis, we repeated the TBS TA-LTP experiments with I_h_ blocked by 20 μM ZD7288. With I_h_ blocked, TBS significantly potentiated TA EPSPs in wild type, but not *fmr1* KO CA1 pyramidal neurons (Supp. Fig. 2). These results suggest that higher dendritic I_h_ alone does not account for the lack of TA-LTP in *fmr1* KO CA1 pyramidal neurons.

### Ca^2+^ entry into *fmr1*KO neurons is reduced at TA synapses

A rise in intracellular Ca^2+^ during TBS is necessary for the induction of TA-LTP^19-21^. We used 2-photon imaging to directly measure changes in dendritic Ca^2+^ during TA TBS in wild type and *fmr1* KO CA1 neurons (Fig. 2a). We varied the initial EPSP amplitude and triggered bursts TA EPSPs (10 at 100 Hz). Small but detectable Ca^2+^ signals were observed at EPSP amplitudes greater than 1 mV for both wild type and *fmr1* KO neurons. In both wild type and *fmr1* KO dendrites, the Ca^2+^ signal increased with increasing EPSP amplitude; however, the Ca^2+^ signal was larger in wild type compared to *fmr1* KO dendrites (Fig. 2b-c).

**Fig. 2:**
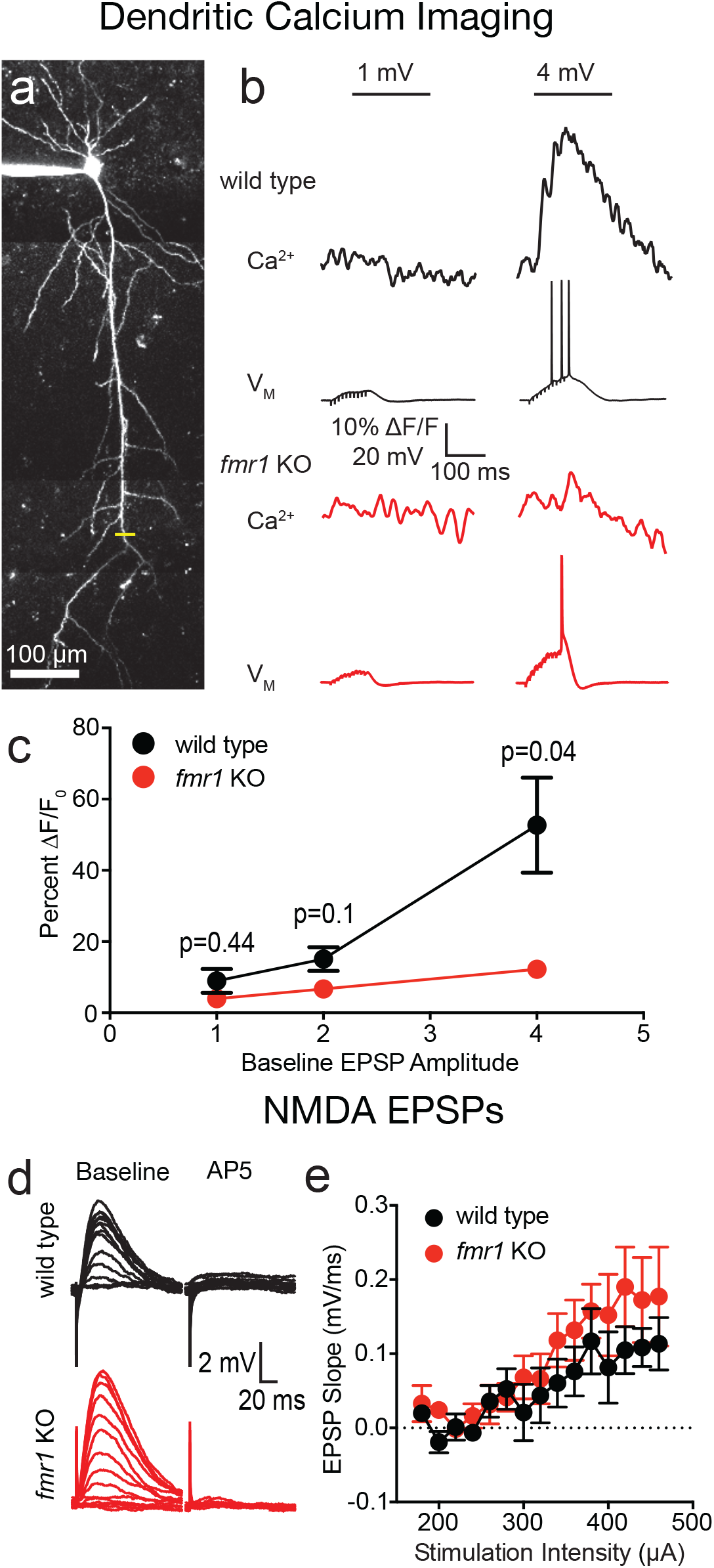
Synaptic Ca^2+^ signaling reduced in fmr1 KO dendrites. **a,** Representative CA1 neuron filled with OGB-1 (100 μM) and Alexa 594 (40 μM). Yellow bar represents location of linescan imaging in the s.l.m region. **b,** Representative Ca^2+^ and voltage signals during 100 Hz bursts of TA EPSPs using 1 and 4 mV initial EPSP amplitudes. **c,** Group data of peak intracellular Ca^2+^ signal during bursts of synaptic activity as a function of initial EPSP amplitude (2-way RM ANOVA: F(1,19)=13.14, p=0.002, η^2^=0.13. Interaction: F(2, 38): 4.86, p=0.013, η^2^=0.1. Sidak’s test-4 mV: p=0.04). **d,** Representative input output traces of NMDAR dependent EPSPs in 0 mM Mg^2+^ and the AMPA blocker DNQX (20 μM) (left). Addition of the NMDAR antagonist AP5 (25 μM) confirmed the synaptic response was NMDAR dependent (right). **e,** Slope of NMDAR EPSPs as a function of stimulation intensity was not different between wild type and *fmr1* KO neurons (2-way RM ANOVA: F(1,12)=0.19, p=0.87).

### NMDAR-mediated EPSPs are not different between wild type and fmr1 KO TA synapses

N-methyl-D-aspartate receptor (NMDAR) activation is a key source of Ca^2+^ influx during the induction of TA-LTP^19-21^. Although we previously showed that there was no significant difference in TA EPSPs between wild type and *fmr1* KO CA1 neurons (Fig. 1), those experiments did not separate the AMPA and NMDAR contributions to the EPSP. To test if NMDAR-mediated EPSPs are different between wildtype and *fmr1* KO neurons, we stimulated TA inputs in the presence of AMPA receptor blocker DNQX (20 µM) and with 0 mM Mg^2+^ in the extracellular saline (Fig. 2d, left). Application of the NMDAR antagonist D-AP5 (50 µM) confirmed isolation of NMDAR-mediated EPSPs in a subset of experiments (Fig. 2d, right). In agreement with our data in Fig. 1e, we found no difference in isolated NMDAR-mediated TA EPSPs between wild type and *fmr1* KO CA1 pyramidal neurons (Fig. 2e).

### Dendritic recordings of TA synaptic transmission and LTP

Using somatic recordings we showed a clear lack of TA-LTP in *fmr1* KO CA1 pyramidal neurons and dendritic imaging revealed reduced dendritic Ca^2+^ influx during bursts of TA stimulation. This suggests that the dendrites are the locus of the changes that impair TA-LTP in *fmr1* KO CA1 pyramidal neurons. Many of the dendritic events required for TA-LTP are distorted or undetectable using somatic recording due to the filtering properties of CA1 dendrites. Thus, we performed current clamp recordings from the apical dendrites of wild type and *fmr1* KO CA1 pyramidal neurons (Fig. 3a; distance from soma–wild type: 211.8 ± 3.52 µm; *fmr1* KO 212.1 ± 4.69 µm; t(28)=0.043; p=0.97). Consistent with our somatic recordings, TA EPSP slope was significantly increased after TBS in wild type but not *fmr1* KO neurons (Fig. 3b-c). There was no difference in the area under the curve during TBS of TA synapses between wild type and *fmr1* KO CA1 pyramidal neurons (Fig. 3d). There was no significant difference in the response to single stimuli (Fig. 3e), paired-pulse ratio (Fig. 3f), or summation (Fig. 3g) between wild type and *fmr1* KO CA1 neurons. Paired-pulse ratios were not different before and after TBS (wild type: 2-way RM ANOVA: F(1,6)=1.89, p=0.22; *fmr1* KO: 2-way RM ANOVA: F(1,6)=0.02, p=0.89). Dendritic recordings, much like in somatic recordings, show a lack of *fmr1* KO TA-LTP. We next used dendritic recording to investigate suprathreshold dendritic events implicated in TA-LTP.

**Fig. 3:**
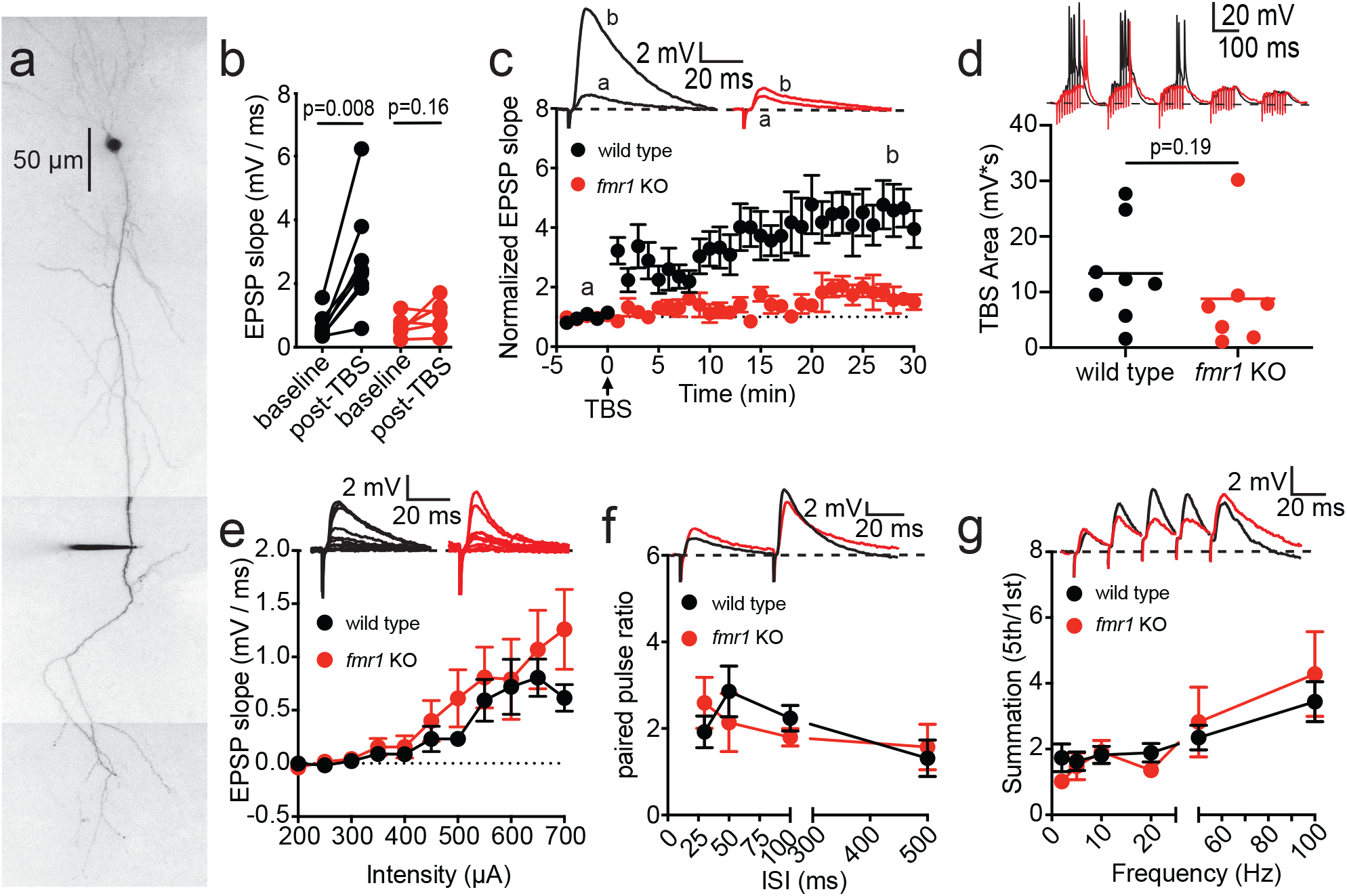
Dendritic recordings show a lack of TA-LTP in fmr1 KO CA1 pyramidal neurons. **a,** Representative dendritic recording from a CA1 neuron filled with Alexa 594 (40 μM). **b,** EPSP slope is increased 30 minutes post-TBS in wild type but not *fmr1* KO CA1 neurons; (wild type: Wilcoxon: W=36, p=0.008, η^2^=0.4; *fmr1* KO: Wilcoxon: W=18, p=0.16). **c,** Normalized EPSP change in EPSP slope for wild type and *fmr1* KO CA1 neurons. *Inset*: Representative EPSP traces from baseline (a) and post-TBS (b). **d,** Graph of area under the curve during TBS (Mann-Whitney: U=16, p=0.19). **e,** Input output of EPSP slope at TA inputs in dendritic recordings (2-way RM ANOVA: F(1,17)=0.63, p=0.44). *Inset*: representative dendritic experiments. **f,** Paired pulse ratio is not different between wild type and *fmr1* KO CA1 pyramidal neurons (2-way RM ANOVA: F(1,12)=0.022, p=0.88). *Inset*: 50 ms ISI paired pulse. **g,** Temporal summation is not different between wild type and *fmr1* KO CA1 pyramidal neurons (2-way RM ANOVA: F(1,12)=0.00001, p=0.99). *Inset*: 50 Hz temporal summation experiment.

### Dendritic complex and Ca^2+^ spikes are not different between wild type and fmr1 KO neurons

NMDARs, activated by presynaptic glutamate release, are one source of Ca^2+^ signaling. We found that subthreshold NMDAR-mediated EPSPs activated by stimulation of TA synapses were not different between wild type and *fmr1* KO neurons. Dendritic voltage-gated channels can also contribute to rises in dendritic Ca^2+19-22^. The activation of distal synapses in CA1 pyramidal neurons gives rise to complex spikes in CA1 dendrites mediated by voltage-gated Na^+^ and Ca^2+^ channels^21,23-26^. We therefore tested the hypothesis that complex spikes and Ca^2+^-dependent action potentials are impaired in *fmr1* KO neurons.

In CA1 pyramidal neurons, dendritic complex spikes consist of a fast initial Na^+^ spike which triggers 1–3 slower, Ca^2+^ mediated spikes^25^. We used dendritic current injection (1 second) to compare complex spikes between wild type and *fmr1* KO CA1 neurons (Fig. 4a). Previous studies using rat hippocampus showed that only a sub-population of CA1 dendrites fire complex spikes^23,25^. We found that approximately half of mouse CA1 neurons fired complex spikes and that the proportion was not different between wild type and *fmr1* KO CA1 pyramidal neurons (wild type: 51.6%, *fmr1* KO: 47.6%). There was no difference in the width of the complex spikes between wild type and *fmr1* KO CA1 dendrites (Fig. 4b-d)^21^. To isolate the dendritic Ca^2+^ spike component of the complex spike, we applied 0.5 µM TTX to block voltage-gated Na^+^ channels and 50 µM 4-AP to block voltage-gated K^+^ channels and bias the dendrites toward firing dendritic Ca^2+^ spikes^23,25,27^. Dendritic Ca^2+^ spikes were evoked by depolarizing current injections and confirmed to be mediated by voltage-gated Ca^2+^ channels by the addition of 200 µM Cd^2+^ (Fig. 4e). The number of elicited Ca^2+^ spikes was not different between wild type and *fmr1* KO CA1 pyramidal neuron dendrites (Fig. 4f). The first Ca^2+^ spike elicited was selected for further analysis (Fig. 4g-k). Dendritic Ca^2+^ spike amplitude (Fig. 4h), maximum rate of rise (Fig. 4i), maximum rate of decay (Fig 4j) and estimated threshold (Fig. 4k) were not different between wild type and *fmr1* KO neurons. These results suggest that impairment of dendritic complex spikes or Ca^2+^ spikes does not contribute to the reduced dendritic Ca^2+^ signal in *fmr1* KO CA1 pyramidal neuron bursts of TA stimulation.

**Fig. 4:**
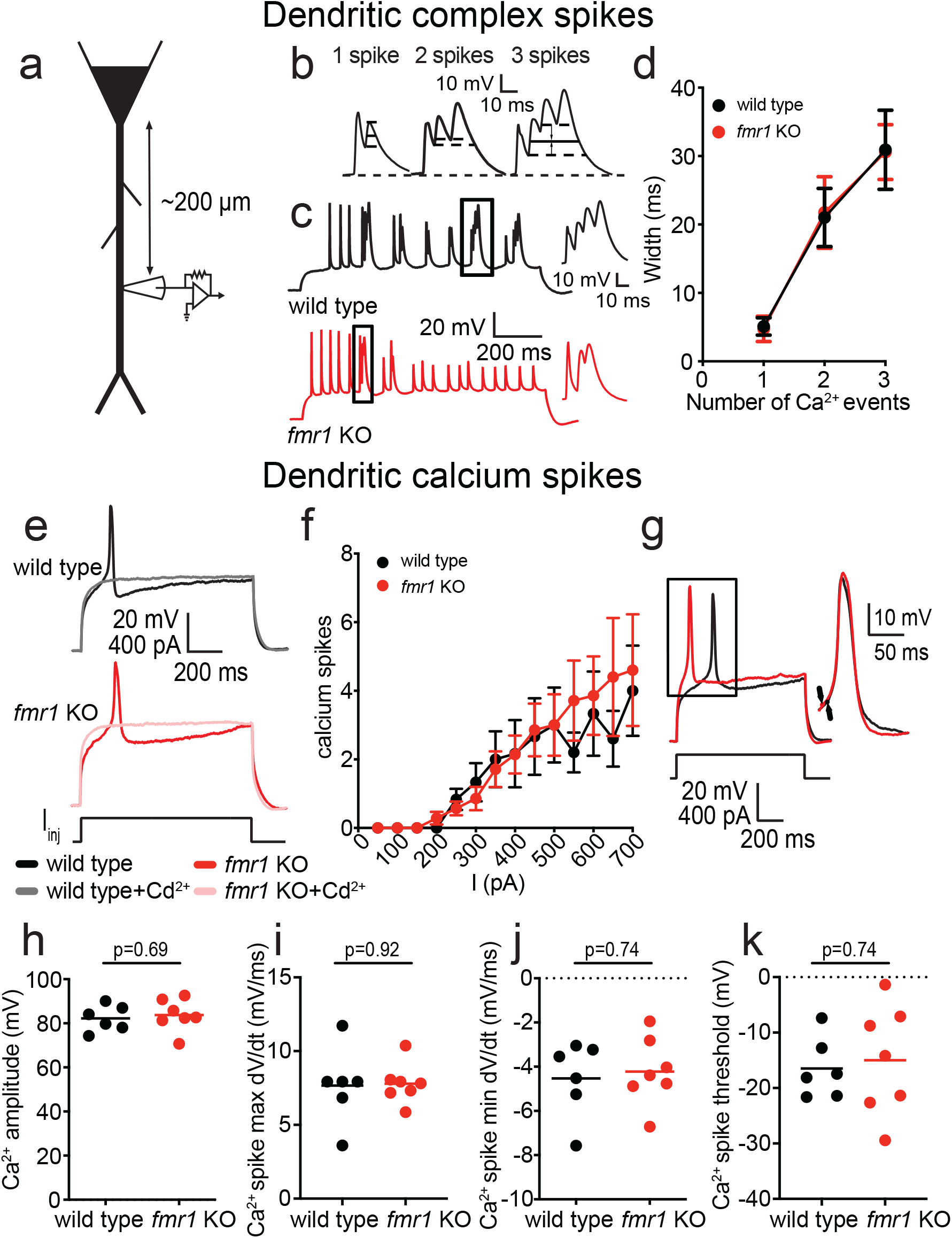
Dendritic complex and Ca^2+^ spikes are not different between wild type and fmr1 KO CA1 pyramidal neurons. **a,** Dendritic recording schematic. **b,** Method for measuring complex spike width (see methods for description). **c,** Complex spikes elicited by dendritic current injection in wild type and *fmr1* KO neurons (left) and example complex spikes (boxes) shown on expanded time scale (right). **d,** Complex spike width as a function of number of events (1-3) (2-way RM ANOVA: F(1,24)=0.00002, p=0.67). **e,** Representative traces of isolated dendritic Ca^2+^ spikes in the presence of 500 nM TTX and 50 µM 4-AP. Application of 200 µM Cd^2+^ confirmed that spikes were generated by voltage gated Ca^2+^ channels. **f,** The number of Ca^2+^ spikes as a function of current amplitude (2-way RM ANOVA: F(1,11)=0.07, p=0.8). **g,** Representative dendritic Ca^2+^ spike traces (left). Single Ca^2+^ spike in the box at left on expanded time scale used for analyses in **h-k** (right). Black arrows show the estimated threshold. **h-k,** Dendritic Ca^2+^ spike amplitude (h, unpaired t-test: t(11)=0.42, p=0.69), maximum rate of rise (i, unpaired t-test: t(7.2)=0.11, p=0.92), minimum rate of decay (j, unpaired t-test: t(11)=0.34, p=0.74), and threshold (k, unpaired t-test: t(9.5)=0.34, p=0.74) were not significantly different between wild type and *fmr1* KO neurons.

### Block of inwardly rectifying K^+^ channels rescues TA-LTP in fmr1 KO neurons

Our results thus far suggest that there are no differences in NMDARs at TA synapses, dendritic complex spikes, or dendritic Ca^2+^ spikes between wild type and *fmr1* KO CA1 pyramidal neurons. Furthermore, despite higher dendritic expression, block of I_h_ in *fmr1* KO neurons does not rescue TA-LTP. Inwardly rectifying K^+^ (K_IR_) channels are open at or near the resting membrane potential, expressed in CA1 pyramidal neuron dendrites and contribute to the induction of hippocampal LTP^28-30^. In rats, K_IR_ channels in the dendrites constrain dendritic nonlinear events and control LTP^29^. To test K_IR_ channels contribute to the lack of TA-LTP in *fmr1* KO neurons, we performed TBS TA-LTP experiments in the presence of 25 μM Ba^2+^ to block K_IR_s ^29^ (Fig. 5a). Blockade of K_IR_ rescued TA-LTP in *fmr1* KO CA1 pyramidal neurons (Fig. 5b-c). The area under the curve during the induction protocol was not different between wild type and *fmr1* KO CA1 pyramidal neurons in the presence of 25 μM Ba^2+^ (Fig. 5d).

**Fig. 5:**
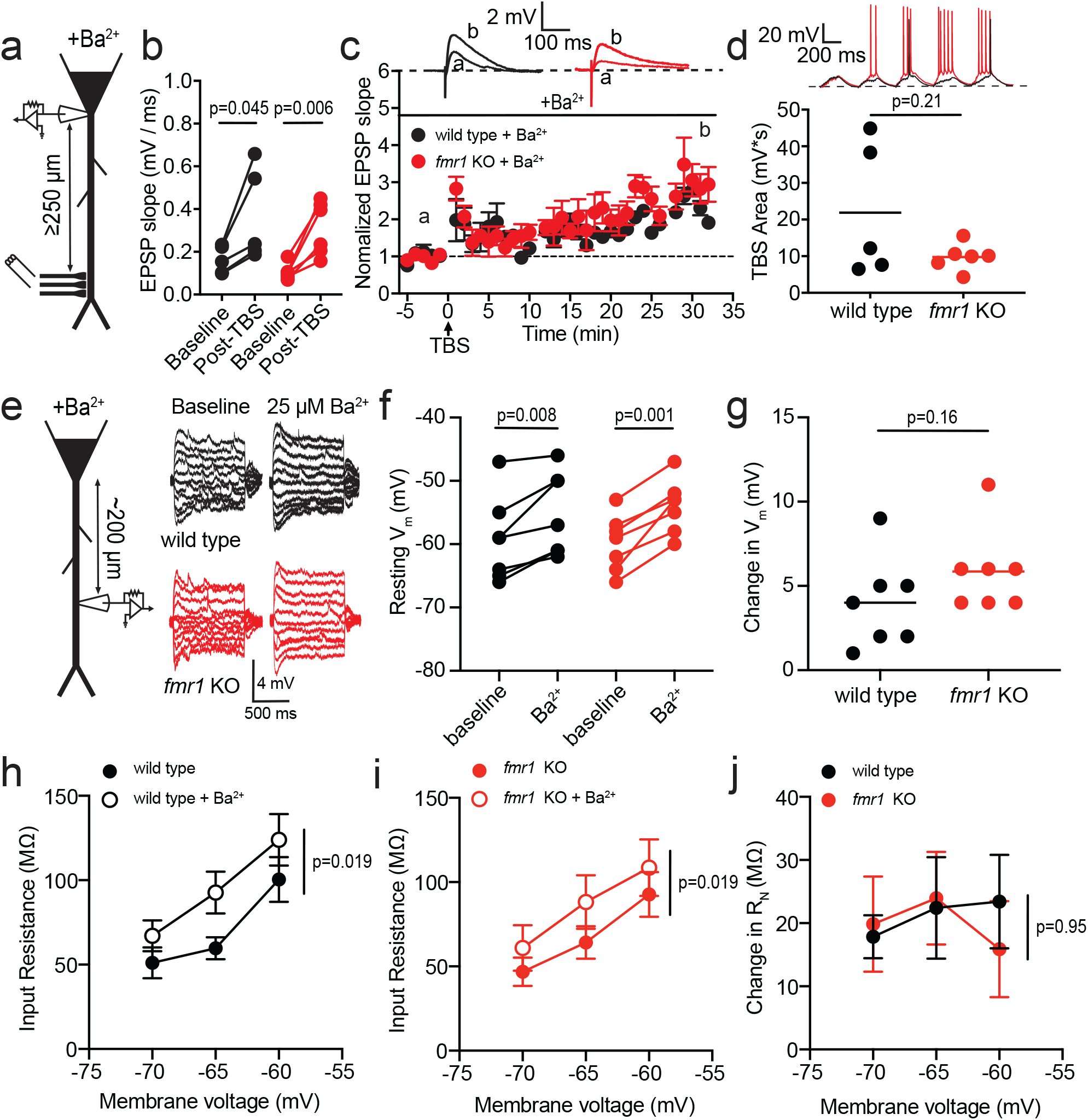
Low Ba^2+^ rescues TA-LTP in fmr1 KO CA1 pyramidal neurons. **a,** Recording configuration during low Ba^2+^ TA-LTP experiments. **b,** EPSP slope is significantly increased 30 minutes after TBS in both wild type and *fmr1* KO CA1 neurons (wild type: paired t-test: t(4)=2.88, p=0.045, η^2^=0.3; *fmr1* KO: paired t-test: t(5)=4.62, p=0.006, η^2^=0.52). **c,** Normalized change in EPSP slope for wild type and *fmr1* KO CA1 neurons in the presence of 25 µM extracellular Ba^2+^. *Inset*: baseline and post-TBS EPSP traces from wild type and *fmr1* KO neurons. **d,** Graph of area under the curve during TBS with Ba^2+^ included in the bath saline (unpaired t-test: t(4.26)=1.46, p=0.21) **e,** Recording configuration (left) and representative voltage traces before and after application of 25 μM Ba^2+^. **f,** Ba^2+^ significantly depolarizes V_M_ in both wild type and *fmr1* KO neurons (wild type: paired t-test: t(6)=3.91, p=0.008, η^2^=0.09; *fmr1* KO: paired t-test: t(6)=6.25, p=0.001, η^2^=0.31). **g,** The effect of Ba^2+^ on V_M_ is not different between wild type and *fmr1* KO neurons (Mann-Whitney: U=13.5, p=0.16). **h, i,** Ba^2+^ significantly increases R_N_ in wild type (h, 2-way RM ANOVA: F(1,5)=11.81, p=0.019, η^2^=0.12) and *fmr1* KO (i, 2-way RM ANOVA: F(1,6)=10.13, p=0.019, η^2^=0.1) dendrites. **j** The effect of Ba^2+^ on R_N_ is not different between wild type and *fmr1* KO neurons (2-way RM ANOVA: F(1,11)=0.005, p=0.95).

One potential explanation for the rescue of TA-LTP by Ba^2+^ is that there is a higher dendritic expression of K_IR_s in *fmr1* KO CA1 pyramidal neurons. We measured the dendritic resting membrane potential (V_m_) and input resistance (R_N_) before and after application of 25 μM Ba^2+^ (Fig. 5e). Extracellular Ba^2+^ depolarized V_M_ in both wild type and *fmr1* KO dendrites (Fig. 5f).

The effect of Ba^2+^ on V_M_ was not significantly different between wild type and *fmr1* KO neurons (Fig. 5g). Extracellular Ba^2+^ also increased R_N_ in both wild type and *fmr1* KO dendrites (Fig. 5h-I). The change in R_N_ was not significantly different between wild type and *fmr1* KO dendrites (Fig. 5j). These results demonstrate that, although block of K_IR_s by extracellular Ba^2+^ rescued TA-LTP in *fmr1* KO neurons, the functional expression of dendritic K_IR_s is not different between wild type and *fmr1* KO CA1 pyramidal neurons.

### Dendritic depolarization rescues TA-LTP in fmr1 KO CA1 pyramidal neurons

Block of K_IR_s depolarized dendritic V_m_, increased dendritic R_N_ and rescued TA-LTP. In rat CA1 neurons, direct manipulation of dendritic V_m_ was able to reproduce the effects of K_IR_s on dendritic function^29^. To test whether the depolarization would mimic the effects of Ba^2+^ on TA-LTP, we depolarized the dendritic V_m_ by 10 mV during the delivery of TBS using steady state current injection to mimic the depolarization of oblique dendrites (see methods) during Ba^2+^ wash-on (Fig. 6a). Similar to extracellular Ba^2+^, dendritic depolarization rescued TA-LTP in *fmr1* KO neurons (Fig. 6b-c). The area under the curve during the induction protocol was not different between wild type and *fmr1* KO CA1 pyramidal neurons (Fig. 6d). Unlike the Ba^2+^ experiments above, however, dendritic V_m_ was only depolarized during delivery of the TBS. Taken together with Ba^2+^ wash-on experiments, these results suggest that *fmr1* KO CA1 dendrites are unable to reach the threshold for TA-LTP induction under normal conditions.

**Fig. 6:**
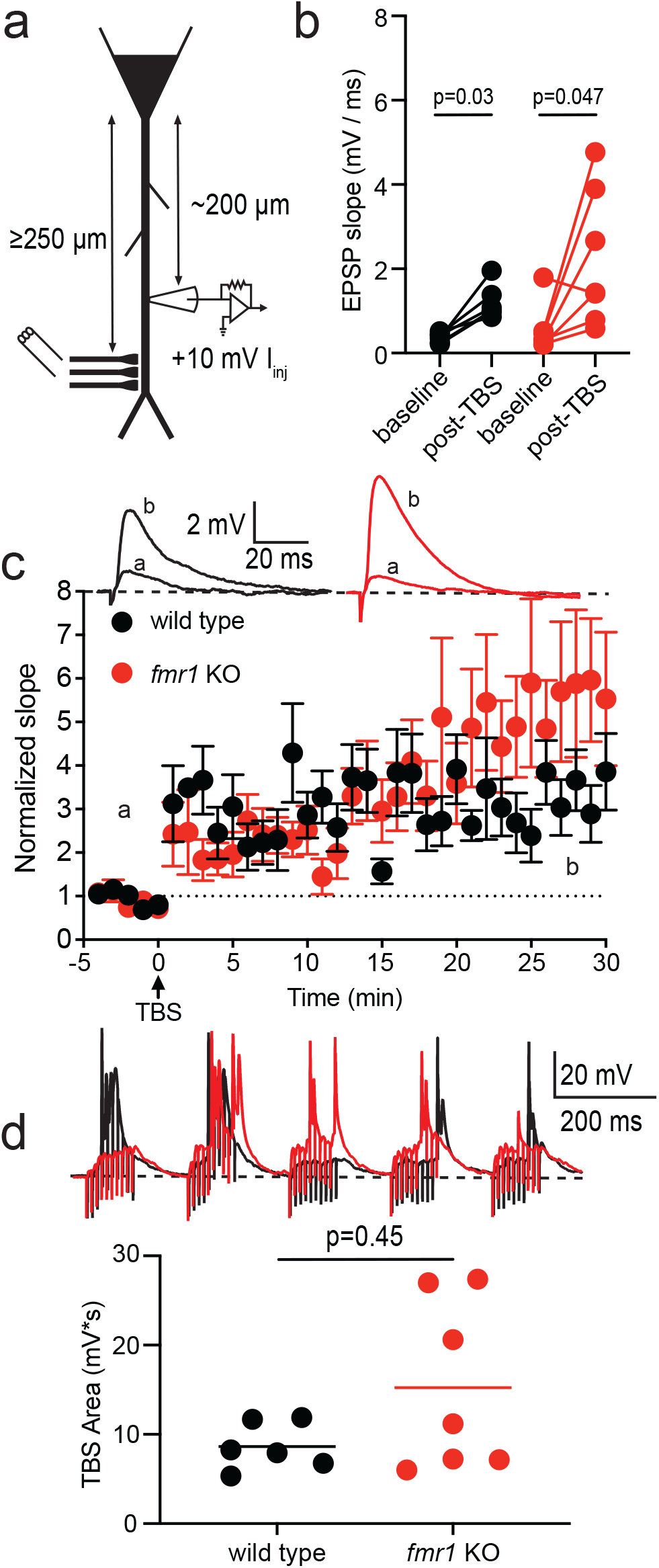
Dendritic depolarization rescues TA-LTP in fmr1 KO CA1 neurons. **a,** Recording configuration. Current injection produced a 10 mV depolarization during TBS. **b,** EPSP slope is significantly increased 30 minutes after TBS in both wild type and *fmr1* KO CA1 neurons (wild type: Wilcoxon: W=21, p=0.03, η^2^=0.4; *fmr1* KO: Wilcoxon: W=24, p=0.047, η^2^=0.52). **c,** Normalized EPSP slope 5 minutes before and 30 minutes after TBS of TA inputs. *Inset*: Representative baseline and post-TBS EPSPs from wild type and *fmr1* KO CA1 neurons. **d,** Graph of area under the curve during TBS while a steady depolarizing current of 10 mV was applied to the dendritic patch (Mann-Whitney: U=15, p=0.45).

### Dendritic spike generation is impaired in fmr1 KO CA1 pyramidal neurons

The large scale, rapid influx of Ca^2+^ necessary for the generation of LTP at distal synapses in CA1 neurons is dependent on dendritic Na^+^ spikes (dspike)^26^. Consequently, one explanation for our results thus far is that dendritic spike generation is impaired in *fmr1* KO neurons. We used a double exponential current injection (τ_rise_ = 0.1 ms; τ_decay_ = 2 ms) to mimic the time course of dendritic EPSPs and trigger dendritic Na^+^ spikes in wild type and *fmr1* KO dendrites (Fig. 7a; see methods). Dspikes in *fmr1* KO CA1 neurons had a more depolarized threshold (Fig. 7b) and slower maximum dV/dt (Fig. 7c) compared to wild type dendrites.

**Fig. 7:**
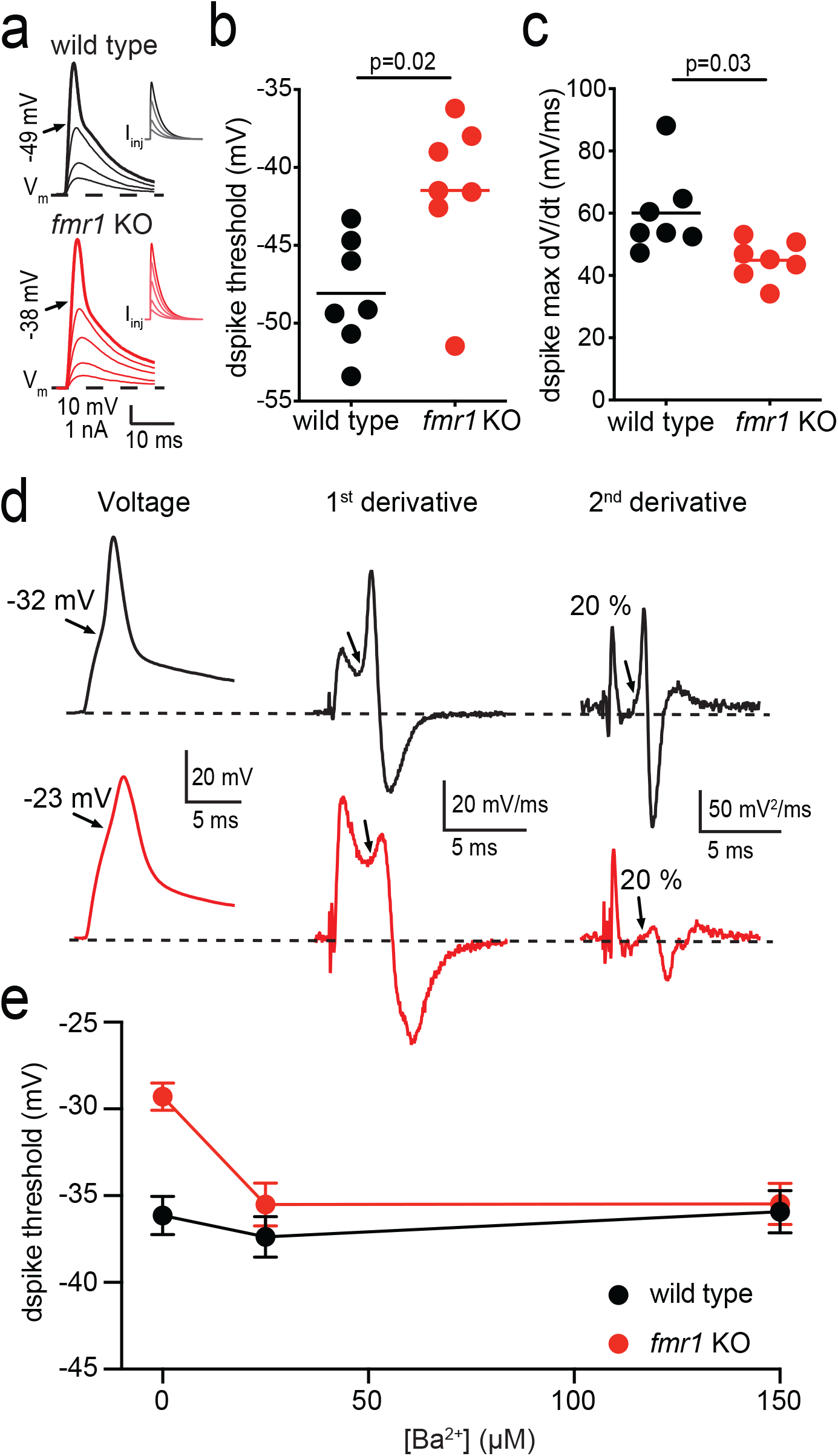
Dendritic spike threshold is more depolarized in in fmr1 KO CA1 pyramidal neurons. **a,** Dendritic voltage response to double exponential current injections (*inset*) of increasing amplitude (500 pA intervals). Thick lines show voltage traces with dspike and the corresponding current injection. **b,** voltage threshold for dspikes is more depolarized in *fmr1* KO CA1 neurons (unpaired t-test: t(10.92)=2.86, p=0.02, η^2^=0.37). **c,** the maximum rate of rise is significantly slower in *fmr1* KO neurons (unpaired t-test: t(8.49)=2.67, p=0.03, η^2^=0.34). **d,** Representative dendritic spike recordings showing recorded voltage (left), 1^st^ derivative (middle), and 2^nd^ derivative (right). Threshold was determined as 20% of the second peak of the second derivative (arrows). **e,** Extracellular Ba^2+^ hyperpolarizes dspike threshold in *fmr1* KO but not wild type dendrites (wild type: n=7 from 3 mice; *fmr1* KO: n=8 from 2 mice; 2-way RM ANOVA: F(1,13)=4.9, p=0.045, η^2^=0.15. Interaction: F(2, 26)=11.51, p=0.0003, η^2^=0.13. Sidak’s test − 0 μM Ba^2+^: p=0.001, η^2^=0.64).

CA1 pyramidal neuron dendrites express voltage-gated Na^+^ channels which contribute to dspikes^22,24^. To test whether dspike threshold and dV/dt differences were caused by differences in Na^+^ channel function we measured Na^+^ current using cell attached voltage clamp recordings from wild type and *fmr1* KO dendrites (200-250 μm from the soma). The maximum current and voltage-dependence of Na^+^ channels we measured in wild type CA1 pyramidal neurons were in good agreement with those obtained from rat CA1 dendrites^22,31^. We found no difference in the Na^+^ I-V curve between wild type and *fmr1* KO neurons (Supplemental Fig. 3a). Additionally, there was no significant difference in the voltage-dependence of activation or steady-state inactivation between wild type and *fmr1* KO Na^+^ channels (Supplemental Fig. 3b-f). Our results suggest that the differences in Na^+^ channel function do not contribute to the threshold and dV/dt differences in dspikes between wild type and *fmr1* KO CA1 pyramidal neurons.

### A-type K^+^ channel block restores dspike threshold and TA-LTP in fmr1 KO neurons

A-type K^+^ currents (I_KA_) are smaller in *fmr1* KO CA1 neurons compared to wild type^15^. We did not initially consider differences in I_KA_ as a potential cause for the lack of TA-LTP in *fmr1* KO neurons as we would expect the reduction in I_KA_ to either promote or have no effect on dspikes. There is, however, a hyperpolarized shift in the activation of A-type K^+^ channels in *fmr1* KO dendrites compared to wild type^15^. A-type K^+^ channels that activate at more negative potentials could overlap with Na^+^ currents and influence the dendritic spike in *fmr1* KO neurons.

Furthermore, low concentrations of Ba^2+^ have been shown to affect transient K^+^ currents expressed in the heart^32^. We thus hypothesized that A-type K^+^ channels influence dendritic spikes in *fmr1* KO but not wild type CA1 neurons. As an initial test of this hypothesis, we recorded dendritic spikes under control conditions, 25 μM Ba^2+^ (used in Fig. 5), and 150 μM Ba^2+^, a concentration known to block A-type K^+^ channels (Fig. 7d)^15^. Ba^2+^ had no effect on dendritic spike threshold in wild type neurons; however, dspike threshold was significantly hyperpolarized by 25 or 150 μM BaCl_2_ in *fmr1* KO neurons (Fig. 7e). These results support the hypothesis that A-type K^+^ channels contribute to the depolarized dspike threshold in *fmr1* KO CA1 neurons. These data also suggest that Ba^2+^ rescued TA-LTP in *fmr1* KO neurons by hyperpolarizing the threshold for dspike generation.

To test if block of A-type K^+^ channels rescued TA-LTP in *fmr1* KO neurons, we repeated the TBS TA-LTP experiments in the presence of the A-type K^+^ channel blocker AmmTx3 (500 nM). Blockade of A-type K^+^ channels with 500 nM AmmTx3 rescued TA-LTP in *fmr1* KO CA1 pyramidal neurons (Fig. 8a-c). The area under the curve during TBS was not different while AmmTx3 was present in the bath (Fig. 8d-e).

**Fig. 8:**
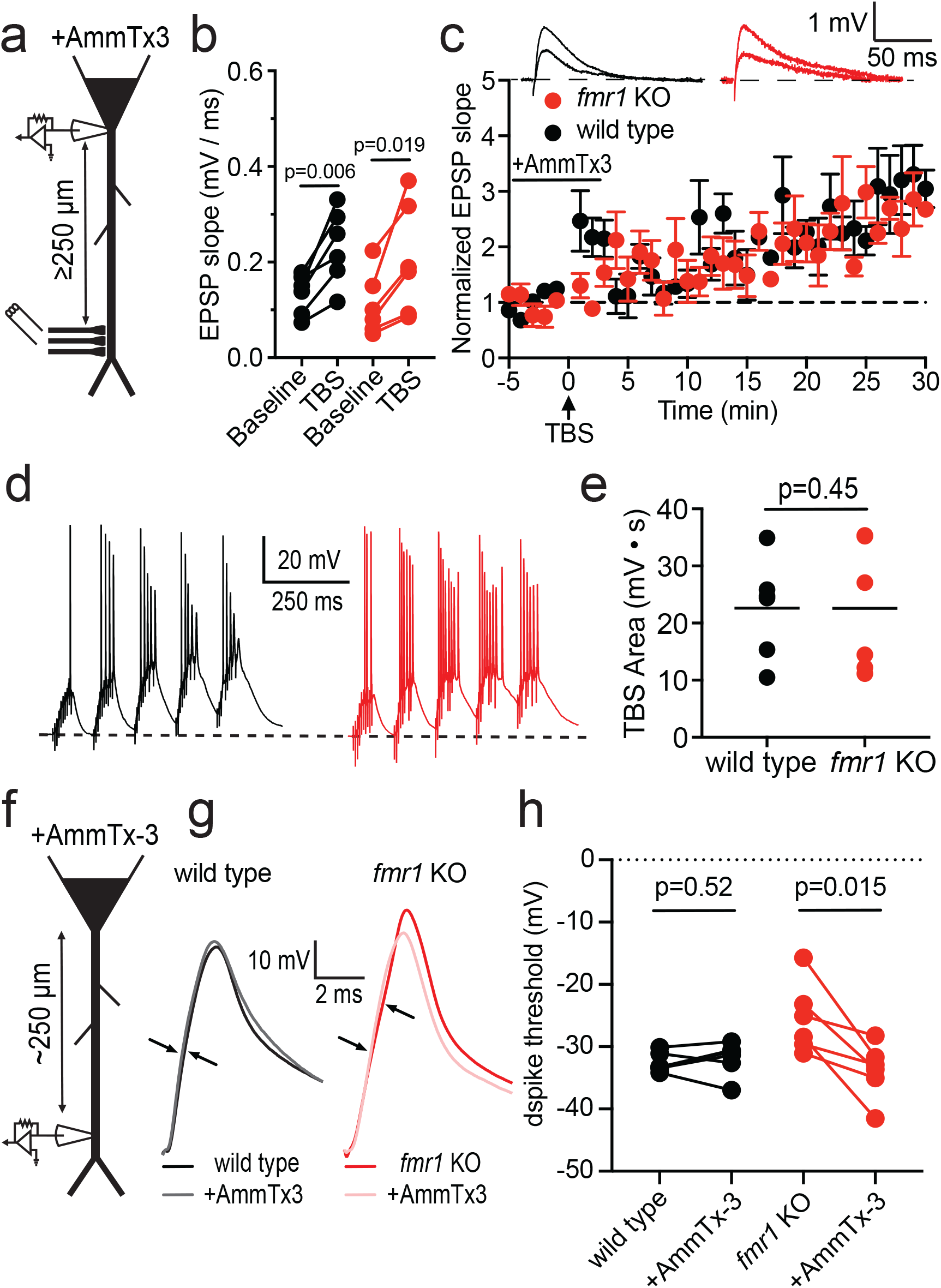
Block of A-type K^+^ channels restores TA-LTP and hyperpolarizes dspike threshold in fmr1 KO neurons. **a,** Recording configuration for AmmTx3 TA-LTP experiments. **b,** EPSP slope is significantly increased 30 minutes after TBS in both wild type and *fmr1* KO CA1 neurons (wild type: n=6 from 3 mice; paired t-test: t(5)=4.6, p=0.006, η^2^=0.39; *fmr1* KO: n=6 from 3 mice; paired t-test: t(5)=3.41, p=0.019, η^2^=0.22). **c,** Normalized EPSP slope 5 minutes before and 30 minutes after TBS of TA inputs. *Inset*: Representative baseline and post-TBS EPSPs from wild type and *fmr1* KO CA1 neurons. **d,** Representative traces showing the somatic response during TBS. **e,** Group data of area under the curve during TBS (unpaired t-test: t(10)=0.008; p=0.99). **f,** Recording configuration for dspike measurements. **g,** Representative dspike recordings from wild type and *fmr1* KO neurons before and after application of 500 nM AmmTx3. Arrows indicate threshold for dspike generation. **h,** AmmTx3 significantly hyperpolarized dspike threshold in *fmr1* KO but not wild type CA1 neurons (wild type: n=6 from 3 mice; paired t-test: t(5)=0.69, p=0.52; *fmr1* KO: n=6 from 3 mice; paired t-test: t(5)=3.64, p=0.015, η^2^=0.41).

We measured the effect of 500 nM AmmTx3 on dspike threshold in wild type and *fmr1* KO CA1 neuron dendrites (Fig. 8f). In agreement with our Ba^2+^ experiments, AmmTx3 significantly reduced dspike threshold in *fmr1* KO CA1 dendrites but had no effect on wild type dspikes (Fig. 8g-h). Taken together, these results suggest that the hyperpolarized shift in activation of A-type K^+^ channels in *fmr1* KO CA1 pyramidal neuron dendrites results in a depolarized threshold for dspikes and a lack of TA-LTP.

## Discussion

Theta-burst stimulation induces behaviorally relevant long-term potentiation of TA inputs in wild type CA1 neurons that is critical to hippocampal-dependent learning and memory^12,13^. We found that TBS of TA inputs fails to induce LTP in *fmr1* KO CA1 pyramidal neurons. We initially hypothesized that the increased functional expression of dendritic h-channels in *fmr1* KO CA1 dendrites would decrease summation of TA inputs and thus impair LTP^5,14^. To our surprise, blocking h-channels with ZD7288 did not rescue TA-LTP in *fmr1* KO neurons. Despite this result, we cannot rule out the possibility that h-channelopathy contributes to difficulty in synaptic stimuli summating to reach dspike threshold and cause TA-LTP induction. LTP at TA synapses requires Ca^2+^ influx through NMDARs and L-type voltage-gated Ca^2+^ channels^19,20^. Two photon Ca^2+^ imaging demonstrated that synaptically evoked dendritic Ca^2+^ signals were consistently smaller in *fmr1* KO neurons. In agreement with a previous study, we found that basal synaptic transmission at TA synapses was not different between wild type and *fmr1* KO mice as has been described previously^10^ (but see^11^). There was no difference in the function of NMDARs, dendritic complex spikes, or isolated dendritic Ca^2+^ spikes between wild type and *fmr1* KO CA1 pyramidal neurons. Dendritic Na^+^ mediated spikes are necessary and act as a trigger for the influx of Ca^2+^ necessary for TA-LTP^19,26^. Our experiments revealed a depolarized dspike threshold that is likely associated with the dysfunction of *fmr1* KO TA-LTP. This suggests that TA-LTP impairment in *fmr1* KO neurons is a result of the inability of TBS depolarization at TA synapses to trigger dspikes. Indeed, depolarization of the dendritic membrane potential in *fmr1* KO neurons through block of inwardly rectifying K^+^ channels or direct dendritic current injection restored TA-LTP. A-type K^+^ channel expression is reduced in the dendrites of *fmr1* KO CA1 pyramidal neurons and the activation shifted to more hyperpolarized potentials^15^. Despite the reduction of A-type K^+^ channel expression, the shift in activation resulted in a depolarized dspike threshold in *fmr1* KO CA1 dendrites, illustrated by the rescue of TA-LTP and dspike threshold in *fmr1* KO neurons by the addition of the A-type K^+^ channel blocker AmmTx3.

### Dendritic nonlinear events in fmr1 KO CA1 neurons

We provide the first direct comparison of dendritic nonlinear events (complex spikes, Ca^2+^ spikes, and dendritic Na^+^ spikes) between wild type and *fmr1* KO CA1 pyramidal neurons. Dendritic complex spikes, which are the combined effect of dendritic Na^+^ and Ca^2+^ channels, were not different in frequency or width between wild type and *fmr1* KO CA1 pyramidal neurons. Recordings of isolated dendritic Ca^2+^ spikes suggest that CA1 dendritic voltage-gated Ca^2+^ channels are not different between wild type and *fmr1* KO neurons. Previous studies have shown that changes in voltage-gated Ca^2+^ channels in *fmr1* KO mice occur in a brain region and cell type specific manner^33-35^. Our results further illustrate that changes in ion channel function and expression in FXS are not conserved across brain regions. Fast, Na^+^-dependent dspikes are essential for the induction TA-LTP^26^. We found that the threshold was more depolarized and the maximum rate of rise slower for d-spikes in *fmr1* KO dendrites compared to wild type. We suggest that the difficulty generating d-spikes contributes to the impaired TA-LTP in *fmr1* KO CA1 neurons.

### A-type K^+^ channels contribute to dendritic spikes in fmr1 KO but not wild type CA1 neurons

The threshold for regenerative events is determined by the complement of ion channels and their relative kinetics^36-38^. We provide the first investigation of dendritic voltage-gated Na^+^ channels in CA1 dendrites of *fmr1* KO mice. The maximum current and biophysical properties of voltage-gated Na^+^ currents in our wild type mouse dendritic recordings were comparable to previous findings in rat^22,31^. Surprisingly, despite the depolarized threshold for dspikes, there was no difference in dendritic Na^+^ channels between wild type and *fmr1* KO CA1 pyramidal neurons. Although block of K_IR_s with low extracellular Ba^2+^ rescued TA-LTP, the effect dendritic V_m_ and R_N_ was not different between wild type and *fmr1* KO CA1 pyramidal neurons, suggesting that dendritic K_IR_s are not different.

We previously found that the activation of A-type K^+^ channels is hyperpolarized in *fmr1* KO CA1 pyramidal neuron dendrites^15^. We hypothesize that the hyperpolarized shift in I_KA_ activation increases the K^+^ conductance during the activation of dendritic Na^+^ channels and depolarizes the threshold for dspike generation. The depolarization produced by extracellular Ba^2+^ could inactivate the A-type K^+^ channels, hyperpolarize dspike threshold, and lead to induction of TA-LTP. Indeed, we found that direct depolarization of the dendrite during TBS was sufficient to rescue TA-LTP in *fmr1* KO CA1 neurons. In support of our hypothesis, both 25 and 150 µM extracellular Ba^2+^ hyperpolarized dendritic spike threshold in *fmr1* KO but not wild type CA1 neurons. We confirmed that block of I_KA_ hyperpolarized dspike threshold and rescued TA-LTP with the specific A-type K^+^ channel blocker AmmTx3.

### Potential mechanism for the hyperpolarized activation of A-type K^+^ channels

The voltage-dependence of A-type K^+^ channels in CA1 pyramidal dendrites is modulated by multiple signaling cascades. Stimulation of protein kinase C (PKC), by activation of metabotropic glutamate and muscarinic acetylcholine receptors, and cAMP-dependent protein kinase A (PKA), by dopaminergic and β-adrenergic receptors, produce a depolarizing shift in the voltage-dependence of activation of A-type K^+^ channels^39,40^.

Reduction in the activity of PKC and/or PKA could account for the hyperpolarizing shift in A-type K^+^ channel activation in *fmr1* KO CA1 neurons. There is evidence from both human and animal models of FXS exhibiting lower cAMP levels^41,42^. Lower cAMP levels would reduce basal PKA activity and potentially lead to a hyperpolarized shift A-type K^+^ channel activation. An early study on cortical synaptoneurosomes found that basal PKC activity was not different between wild type and *fmr1* KO mice^43^. A more recent study, however, found that PKCε expression was significant lower in the hippocampus of *fmr1* KO mice^44^. While not a canonical PKC isoform, in that it is not sensitive to intracellular Ca^2+^, PKCε is abundant in the nervous system and activated by G-protein coupled receptors^45^. Furthermore, PKCε is activated by phorbol esters, which were used to investigate PKC-dependent modulation of A-type K^+^ channels^39^. Thus, PKCε is a potential candidate for regulating the voltage-dependence of A-type K^+^ channel activation. Additionally, FMRP is positive regulator of both PKA^46^ and PKC^47^ and therefore loss of FMRP in FXS could result in lower overall PKA and PKC activity. This suggests that changes in the basal activity of PKA and/or PKC could contribute to the hyperpolarized shift in A-type K^+^ channel activation that alters dendritic spike threshold and impairs TA-LTP in *fmr1* KO CA1 neurons.

### Consequences for hippocampal learning and memory

There are two major changes in A-type K^+^ channel function in *fmr1* KO CA1 dendrites: lower maximum current and hyperpolarized activation. These changes in dendritic A-type K^+^ channels have different effects on LTP at the two major excitatory inputs to CA1 pyramidal neurons. At the proximal and peri-somatic Schaffer collateral inputs, the decreased expression of A-type K^+^ channels increases back-propagating action potential amplitude and resultant Ca^2+^ influx which subsequently reduces the threshold for theta burst pairing LTP in *fmr1* KO neurons^15^. The lower dendritic I_KA_ should also favor dendritic spike initiation. We suggest however, that the hyperpolarized shift in activation antagonizes dendritic Na^+^ channels resulting in a depolarized dspike threshold and prevents the induction of TA-LTP in *fmr1* KO CA1 pyramidal neurons. The coordinated plasticity of TA inputs, which convey current information about the external environment, and Schaffer collateral inputs, which convey stored information from prior experiences, is critical for hippocampal dependent memory tasks. The combined changes in A-type K^+^ channel function alter the dendritic processing of these two critical excitatory pathways. As a result of these changes in LTP, CA1 neurons may have difficulty in establishing and maintaining place fields that would allow for the completion of spatial memory dependent tasks^13^. Indeed, recent findings using in vivo electrophysiological recordings from *fmr1* KO mice identified impairments in hippocampal dependent learning tasks associated with reorienting to changes in rules surrounding the task^48^. Our findings show an inability to induce LTP in TA synapses, in a pathway critical for the induction of long-term memory. This finding provides critical new information for the understanding of the FXS disease phenotype.

FXS is marked by deficits in learning and memory, making the discovery of specific therapeutic targets necessary for the understanding and future treatment of the disorder. Our previously published work, along with other studies, showed how changes in the expression of A-type K^+^ channels affect Schaffer collateral LTP in FXS^15,49,50^. Here, we provide evidence that changes in the biophysical properties of A-type K^+^ channels impact TA-LTP. This suggests that while A-type K^+^ channels represent a potential therapeutic target for FXS interventions, manipulations must take into account both the change in overall expression and the shift in gating.

## Methods

### Animals

The University of Texas at Austin Institutional Animal Care and Use Committee approved all animal procedures. Male wild type and *fmr1* KO C57/B6 mice from 8-16 weeks were used for all experiments. *fmr1* KO male and homozygous female *fmr1* KO mice were paired to produce litters of *fmr1* KO animals. Mice were weaned at P20. Animals were housed in single sex groups at room temperature with *ad libitum* access to food and water and set on a reverse 12-hour light cycle in the University of Texas at Austin vivarium located in the Norman Hackerman Building.

### Preparation of acute brain slices

Mice were anesthetized with acute isofluorane exposure followed by injection of ketamine/xylazine cocktail (100/10 mg/kg i.p.). Mice were then perfused through the heart with ice-cold saline consisting of (in mM): 2.5 KCl, 1.25 NaH_2_PO_4_, 25 NaHCO_3_, 0.5 CaCl_2_, 7 MgCl_2_, 7 dextrose, 205 sucrose, 1.3 ascorbate and 3 Na^+^ pyruvate (bubbled with 95% O_2_/5% CO_2_ to maintain pH at ∼7.4). The brain was removed, trimmed, and sectioned into 300 μm thick transverse slices of middle hippocampus using a vibrating tissue slicer (Vibratome 3000, Vibratome Inc.)^5,14^. Slices were held for 30 minutes at 35°C in a chamber filled with artificial cerebral spinal fluid (aCSF) consisting of (in mM): 125 NaCl, 2.5 KCl, 1.25 NaH_2_PO4, 25 NaHCO_3_, 2 CaCl_2_, 2 MgCl_2_, 10 dextrose and 3 Na^+^ pyruvate (bubbled with 95% O_2_/5% CO_2_) and then at room temperature until the time of recording.

### Electrophysiology

Slices were submerged in a heated (32–34°C) recording chamber and continually perfused (1−2 mL/minute) with bubbled aCSF containing (in mM): 125 NaCl, 3.0 KCl, 1.25 NaH_2_PO_4_, 25 NaHCO_3_, 2 CaCl_2_, 1 MgCl_2_, 10 dextrose, 3 Na^+^ pyruvate, 0.005 CGP and 0.002 gabazine. To reduce recurrent excitation, a cut was made between area CA3 and area CA1. CA1 pyramidal neuron dendrites or somata were visually identified using DIC or Dodt contrast optics.

### Whole-cell current clamp recordings

Patch pipettes (4−8 MΩ somatic, 7–11 MΩ dendritic) were pulled from borosilicate glass and filled with (in mM): 120 K-Gluconate, 16 KCl, 10 HEPES, 8 NaCl, 7 K_2_ phosphocreatine, 0.3 Na−GTP, 4 Mg−ATP (pH 7.3 with KOH). Neurobiotin (Vector Laboratories; 2%) was included in the internal recording solution to determine the recording location during post-hoc morphological reconstruction. Neurons that had a significant portion of the oblique or apical dendrites cut were excluded from analysis. In some cases, Alexa 594 was used to provide real time feedback of dendritic morphology.

Data were acquired using a Dagan BVC−700 amplifier and Axograph (Axograph X) or custom data acquisition software written using Igor Pro (Wavemetrics). Data were sampled at 10−50 kHz, filtered at 3-5 kHz and digitized using an ITC-18 interface (InstruTech). Pipette capacitance and series resistance were monitored and adjusted throughout each recording. Series resistance was monitored throughout each experiment and the experiment discarded if series resistance exceeded 30 MΩ (50 MΩ for dendritic recordings). Experiments in which the resting membrane potential was more depolarized than −50 mV were discarded. The liquid junction potential was estimated to be 14.3 mV (Patcher’s Power tools IGOR Pro) and was not corrected for.

Extracellular stimulation was performed using bipolar sharp tungsten electrodes (5 MΩ, 8° taper, A-M Systems) connected to a Neurolog NL800A current stimulus isolator (Digitimer). The temporoammonic inputs were targeted by visually locating the axon fibers in the SLM region (≥250 μm from CA1 *stratum pyramidale*) and lowering the tungsten electrode until the tip was approximately 10 μm below the surface of the tissue. Stimulation intensity was increased until reliable EPSP (1-2 mV for somatic recordings and 2-4 mV for dendritic recordings) was elicited. To isolate NMDAR dependent EPSPs (Fig. 2) MgCl_2_ was removed from the extracellular aCSF and 20 µM DNQX added to block AMPA receptors.

### Induction of long-term potentiation

Long-term potentiation was performed using theta burst stimulation as previously described^16^. Baseline EPSPs were stimulated at 0.067 Hz for 5 minutes. LTP was induced using TBS with bursts of 10 stimuli at 100 Hz, performed in 5 trains at 5 Hz, and each set of TBS was repeated 4 times at 20 second intervals. EPSPs were then recorded for approximately 30 minutes post TBS. The change in EPSP slope was plotted normalized to the baseline period. In experiments when dendritic depolarization during TBS was applied cells were depolarized by 10 mV using steady state current injection to account for decay of injected current throughout the dendritic arbor^51^. AmmTx3 was applied during TBS at a concentration of 500 nM based on previously published results^52^.

### Analysis of current clamp data

Data were analyzed using Axograph X or custom analysis software written in Igor Pro. EPSP summation was quantified as the ratio of the amplitude of the fifth EPSP to the amplitude of the first EPSP. Paired pulse ratio was calculated as the slope of the second EPSP divided by the slope of the first EPSP. Input-output was measured by increasing the magnitude of stimulus and measuring the slope of the resulting EPSP. For a given input-output experiment (Figs. 1-3) the same stimulation electrode was used throughout to reduce variability within the experiment (but note that stimulation amplitudes differed across the experiments).

Complex spikes were elicited by delivering square current injections (50-450 pA for 1 second). Complex spikes consist of a fast initial spike followed by 1 to 3 slower Ca^2+^ dependent spikes and were categorized as described previously^25^. The width of a complex spike was calculated differentially depending on the number of Ca^2+^ dependent events following the initial fast spike. For one slow spike the width was calculated as the half-width from the initiation point of the spike to the peak of the event. When two or three Ca^2+^ dependent spikes occurred, the width was calculated using methods for the measurement of dendritic Ca^2+^ plateau width modified from^21^. For these events, width was taken as the halfway point between the initiation point of the first slow event and the lowest trough between events (illustrated in Fig. 4b).

Dendritic Ca^2+^ spikes were isolated with the addition of 0.5 μM TTX and 50 μM 4-AP. Spikes were generated by injecting square current pulses 1000 ms long and varying between 50 and 700 pA with 50 pA intervals. Because Ca^2+^ dependent spikes have a much slower time course compared with Na^+^ dependent spikes, measurements that utilized the second derivative of the voltage response (see below) are not reliable. The threshold of somatically recorded action potentials occurs at approximately 8% of the maximum dV/dt. We therefore estimated the Ca^2+^ spike threshold using the time when dV/dt was 8% of the maximum dv/dt. Amplitude was measured as the difference from baseline membrane potential to the peak of the first Ca^2+^ spike.

Dendritic Na^+^ spike experiments were performed on dendritic recordings between 200 and 300 μm from the soma. Spikes were generated by injecting a series of double exponential currents (τ_1_ = 0.1 ms τ_2_ = 2 ms) ranging in amplitude between 500 pA and 5000 pA at 100 or 500 pA intervals. The threshold for dspikes was calculated as 20% of the second peak of the second derivative of the voltage response^53^.

R_N_ was calculated from the linear portion of the current−voltage relationship generated in response to a family of current injections (−50 to +50 pA, 10 pA steps).

### 2-photon Ca^2+^ imaging

Ca^2+^ imaging experiments are performed on a Prairie Ultima 2-photon imaging system (now Bruker) arranged for *in vitro* patch clamp recording. An ultra fast, pulsed laser beam (Spectra Physics: MaiTai) was used at 920 nm for imaging. Recording pipettes were filled with OGB-1 (Invitrogen, 100 μM) and Alexa 594 (Invitrogen, 40 μM). Line scans across the distal dendrites were performed at 500 Hz with a dwell time of 4 μs for between 400 and 1200 ms. Imaging location was chosen approximately 50 μm more proximal to the soma in reference to the extracellular stimulating electrode.

Analysis of line scans were performed using ImageJ (NIH) by placing a line through the fluorescent signal and plotting the profile before converting to a number array^54^. Changes in Ca^2+^ were quantified using ΔF/F, where F is the baseline fluorescence prior to stimulation and ΔF is the change in fluorescence during neuronal stimulation. Ca^2+^ signal traces were smoothed using a Savitsky-Golay function (IGOR pro, Wavemetrics).

### Voltage Clamp Recordings of Na^+^ current

Cell attached voltage clamp recordings were performed using an Axopatch 200B amplifier (Molecular Devices). Data were acquired at 50 kHz and filtered at 2 kHz and then digitized using an ITC-18 interface (InstruTech) and recorded using custom Igor Pro software (Igor Pro 7, Wavemetrics). Pipette solution for cell attached Na^+^ channel recordings consisted of (in mM): 140 NaCl, 2.5 KCl, 2 CaCl_2_, 1 MgCl_2_, 10 HEPES, 10 TEA, 1 3,4 D-AP, 1 4-AP. Na^+^ currents were elicited using depolarizing voltage commands from −70 to 20 mV in 10 mV steps from a holding potential of −90 mV. Steady state inactivation was measured using depolarizing test pulses to a fixed potential (0 mV) preceded by a series of prepulse conditioning potentials ranging from −100 to −20 mV in 10 mV increments. Linear leakage and capacitive currents were digitally subtracted by scaling traces at smaller voltages in which not voltage dependent current was activated. Activation data were plotted as normalized conductance and steady-state inactivation as normalized current. Activation and inactivation data were fit to a single Boltzmann function using a least-squares program.

### DAB reaction and cellular reconstruction

Slices with cells filled with Neurobiotin (Vector Laboratories, Burlingame, CA) during current clamp experiments were fixed in 3% glutaraldehyde for a minimum of 24 hours. Slices were washed in 0.1 M phosphate buffer (PB) and incubated in 0.5% H_2_O_2_ for 30 minutes. Slices were then washed in PB and incubated in ABC reagent (Vector Laboratories, Burlingame, CA) containing Avidin DH and biotinylated horseradish peroxidase H for 24-48 hours at 4°C. Slices were then incubated in DAB solution (Vector Laboratories, Burlingaxme, CA) in the presence of H_2_O_2_ and monitored for a visible color change to the slices and Neurobiotin filled cell. Slices were dehydrated in glycerol and mounted on glass slides for imaging. Identifiable neurons were reconstructed at 40X magnification using a Leitz Diaplan microscope with Neurolucida 6.0 software (MicroBrightField, Inc., Williston, VT). The Sholl radius was set so that there were 10 and 25 concentric circles measuring the basal dendrites and apical dendrites respectively.

### Experimental Design and Statistical Analysis

The use of wild type and *fmr1* KO mice was interleaved during each set of experiments. Where possible, experimenters were blind to the condition of the animal during experimentation and analysis. Experiments were designed as a comparison between wild type and *fmr1* KO mice.

Data were tested for normality using the Shapiro-Wilk normality test and analyzed using an unpaired t-test, paired t-test, a Mann-Whitney rank sum test, or Wilcoxon matched pairs (for paired, non-normal data). Two-way repeated measures (RM) ANOVA was applied to experiments with multiple test variables for each genotype. Sidak’s multiple comparison test was used to compare row means between groups. Data are presented as mean ± standard error. Alpha was set to 0.05 for all experiments. Effect size, a measure of the amount of variance accounted for by differences between groups, is reported as η^2^ only when p<0.05. η^2^ effect sizes are defined as small – 0.01, medium – 0.06, and large – 0.14. Effect sizes from t-tests were initially calculated as Cohen’s d and then converted to η^2^ ^55^.

## Acknowledgments

We thank Dr. Richard Gray for assistance with analysis programs, Meagan Volquardsen for genotyping and mouse colony management, and members of the Johnston lab for helpful comments on this manuscript. This work was supported by National Institutes of Health grant R01 MH100510 (DHB), the Franklyn Alexander Endowed Fellowship (GJO), and an Institutional Training Grant from National Institutes of Health 5T32DA018926.

## Author Contributions

D.H.B. conceived research; G.J.O. and D.H.B. designed experiments; G.J.O., C.J.A., R.A.C. and D.H.B. acquired data; G.J.O. and D.H.B. analyzed data; G.J.O. and D.H.B. interpreted results of experiments; G.J.O. and D.H.B. prepared figures; G.J.O. and D.H.B. drafted manuscript; G.J.O., C.J.A., R.A.C., and D.H.B. edited and revised manuscript.

## Conflict of interest statement

The authors declare no competing interests.

## Notes

### Competing Interest Statement

The authors have declared no competing interest.

