## Supplemental figures for "Altered A-type potassium channel function impairs dendritic spike initiation and temporammonic long-term potentiation in Fragile X syndrome"

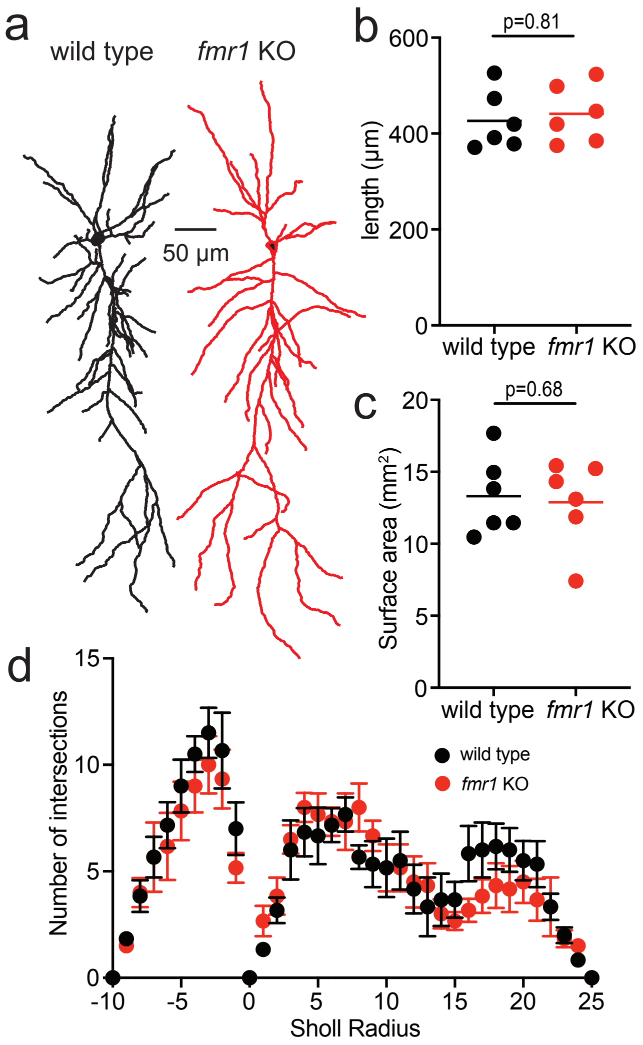


**Supp. Fig. 1: CA1 pyramidal neuron morphology is not different between wild type and *fmr1* KO mice. a,** Representative neuronal reconstructions of wild type (black) and fmr1 KO (red) CA1 pyramidal neurons. **b,c,** Total dendritic length (b, unpaired t-test: t(10)=0.42, p=0.68) and surface area (c, unpaired t-test: t(9.91)=0.25, p=0.81) are not different between wild type and *fmr1* KO CA1 neurons. **d**, Dendritic branching is not different between wild type and *fmr1* KO CA1 neurons (2-way RM ANOVA: F(1,10)=0.19, p=0.67).

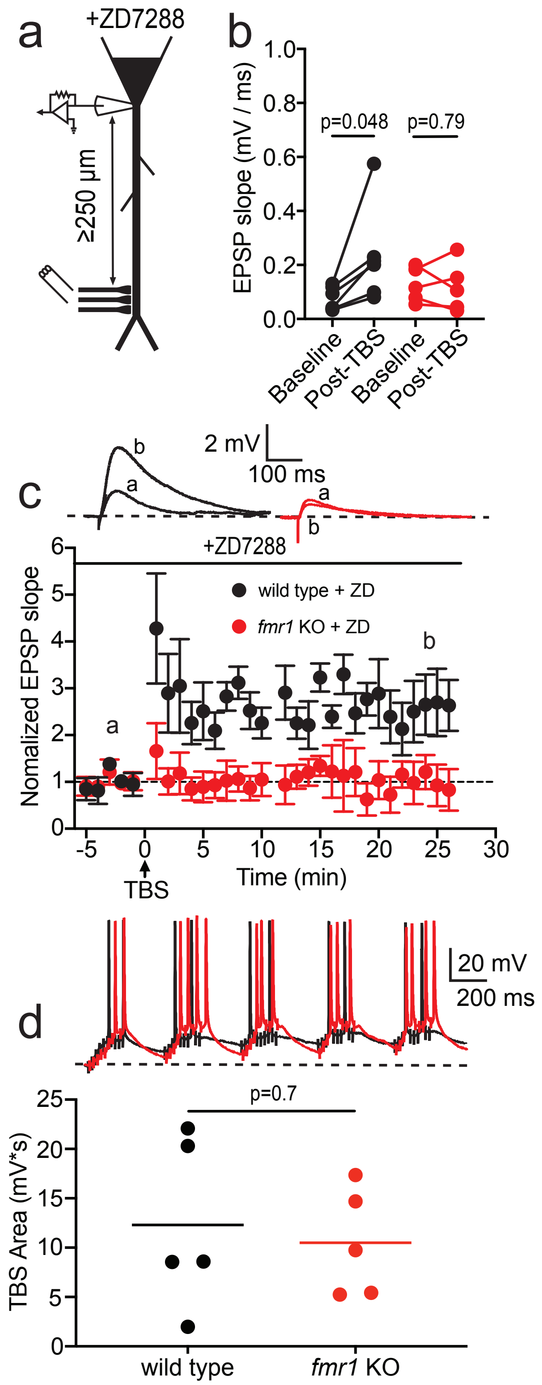


**Supp. Fig. 2: Block of h-channels by 20 µM ZD7288 does not rescue TA-LTP in *fmr1* KO neurons. a,** Recording paradigm during TA-LTP experiments. **b,** EPSP slope is significantly increased 30 minutes after TBS in wild type but not *fmr1* KO CA1 neurons (wild type: paired t-test: $t(5)=2.61$, $p=0.048$, $\eta^2=0.25$. *fmr1* KO: paired t-test: $t(4)=0.28$, $p=0.79$). **c,** Normalized EPSP slope 5 minutes before and 30 minutes after TBS of TA inputs. Inset: Representative baseline and post-TBS EPSPs from wild type and *fmr1* KO CA1 neurons. **d,** Graph of the area under the curve during TBS in the presence of ZD7288 (unpaired t-test: $t(6.7)=0.4$, $p=0.7$).

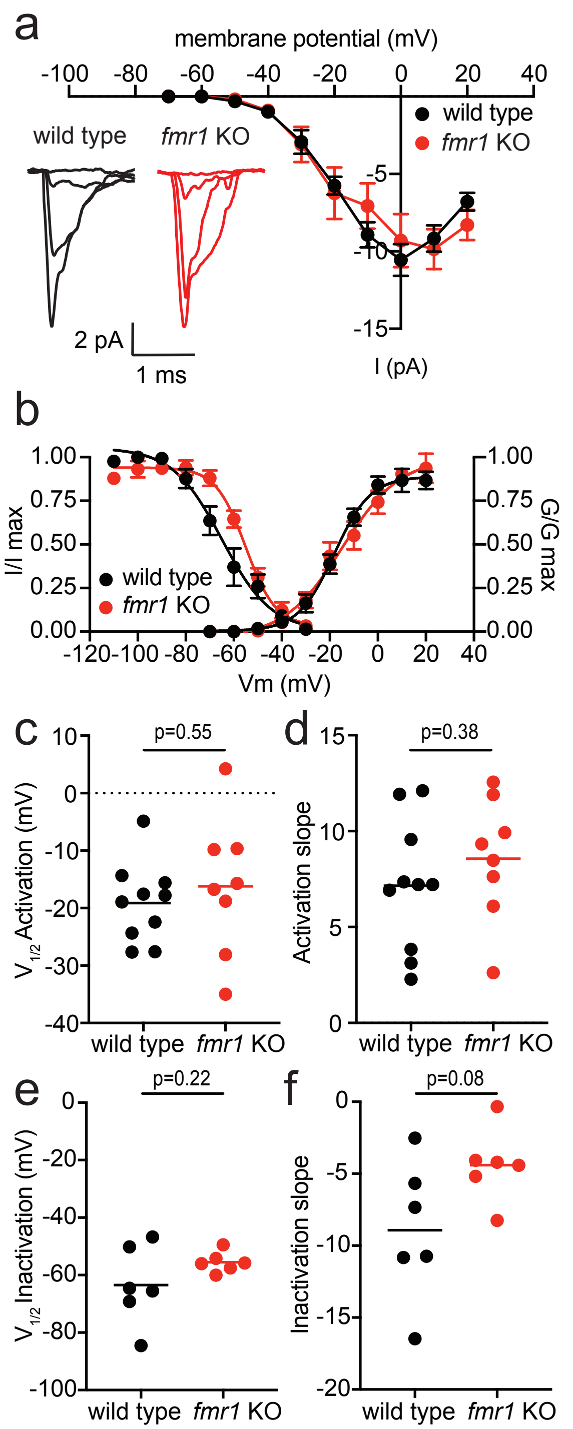


**Supp. Fig. 3: Dendritic Na+ channels are not different between wild type and *fmr1* KO CA1 pyramidal neurons. a,** Current-voltage plot showing no significant difference in Na+ current between wild type and *fmr1* KO dendrites. Inset: representative Na+ current traces in response to voltage steps to -60, -40, -20, and 0 mV. **b,** Na+ channel activation and inactivation are not different between wild type and *fmr1* KO CA1 dendrites. **c,d,** Na+ channel activation V1/2 (c, unpaired t-test: t(10)=0.61, p=0.55) and slope factor (d, unpaired t-test: t(15)=0.9, p=0.38) are not different between wild type and *fmr1* KO dendrites. **e,f** Na+ channel steady state inactivation V1/2 (e, unpaired t-test: t(5.67)=1.38, p=0.22) and slope factor (f, unpaired t-test: t(7.53)=2.02, p=0.08) are not different between wild type and *fmr1* KO dendrites.
